# Tuning retractable, microscale, membrane-breaking protein needles

**DOI:** 10.1101/029587

**Authors:** Jessica K. Polka, Pamela A. Silver

## Abstract

The refractile (R) bodies found in *Caedibacter taeniospiralis,* a bacterial endosymbiont of *Paramecium tetraurelia*, are large, polymeric protein structures that can switch between two conformations. At cytoplasmic pH, they resemble coiled ribbons of protein 500nm in diameter. At low pH, they extend to form hollow needles up to 20 microns long. They can be expressed heterologously from an operon containing four short open reading frames and can function *in vitro* in diverse buffer conditions.

In this study, R bodies purified from *Escherichia coli* were found to be capable of undergoing many consecutive extension-contraction cycles. Furthermore, the solubility of R bodies, which can easily be interpreted by eye, was found to correlate with their extension state. This macroscopic phenotype was used to develop a quantitative, high-throughput assay for R body state, enabling a visual screen of R body mutants defective in extension. The role of specific amino acids in extension was determined, and this information was used to construct rationally-designed mutants tailored to extend at higher pH. Furthermore, R bodies were able to rupture *E. coli* spheroplasts to release soluble proteins across lipid bilayers. Taken together, these results show that R bodies act as tunable, pH-actuated pistons suitable for a variety of membrane-breaking applications.

## Importance

R bodies are natural toxin delivery machines made by bacteria that live inside of single-celled eukaryotes. Under normal conditions, they resemble large, coiled protein ribbons. However, under acidic conditions (such as those encountered when ingested by a eukaryotic cell), they dramatically extend into a long, hollow tube that can disrupt membranes. R bodies are made from only four small proteins, function independently of cells, and can withstand harsh conditions. As such, they hold promise as tools to facilitate gene or drug delivery.

Here we show the R body extension process is reversible over many cycles and that R bodies are capable of releasing *E. coli* cell contents into the environment. Furthermore, we generated a panel of mutant R bodies that extend at varying pH values. These mutants demonstrate that R bodies can be tuned to function in specific applications.

## Introduction

R bodies (Type 51 refractile bodies) are ribbon-like protein polymers that are naturally expressed in the cytoplasm of *Caedibacter taeniospiralis*, an endosymbiont of “killer” strains of *Paramecium tetraurelia* (reviewed in (1)). These bacteria, also called kappa particles, confer to their host the ability to kill other strains of *Paramecium.* This killing is dependent on ingestion of the R body-containing bacteria (2) that are shed into the environment by the killer strain (3, 4). Inside the food vacuole of the non-killer paramecium, acidic conditions cause the R body to unroll from a coil 500nm in diameter to form a tube 165nm in diameter and up to 20μm long (Figure 1A-C). This extension deforms and punctures the membrane of the food vacuole, mixing contents of the bacteria with the paramecium’s cytoplasm (2, 5). The subsequent death of the paramecium presumably results from the release of unidentified toxins from the bacteria into the cytoplasm, as killing does not occur when sensitive strains are fed purified R bodies or *E. coli* expressing R bodies (6–8). Thus, R bodies themselves are not lethal, but rather they have been proposed to act as delivery devices (Figure 1A). This process confers a competitive advantage to the killer paramecium (9) and therefore benefits its endosymbionts as well.

**Figure 1:**
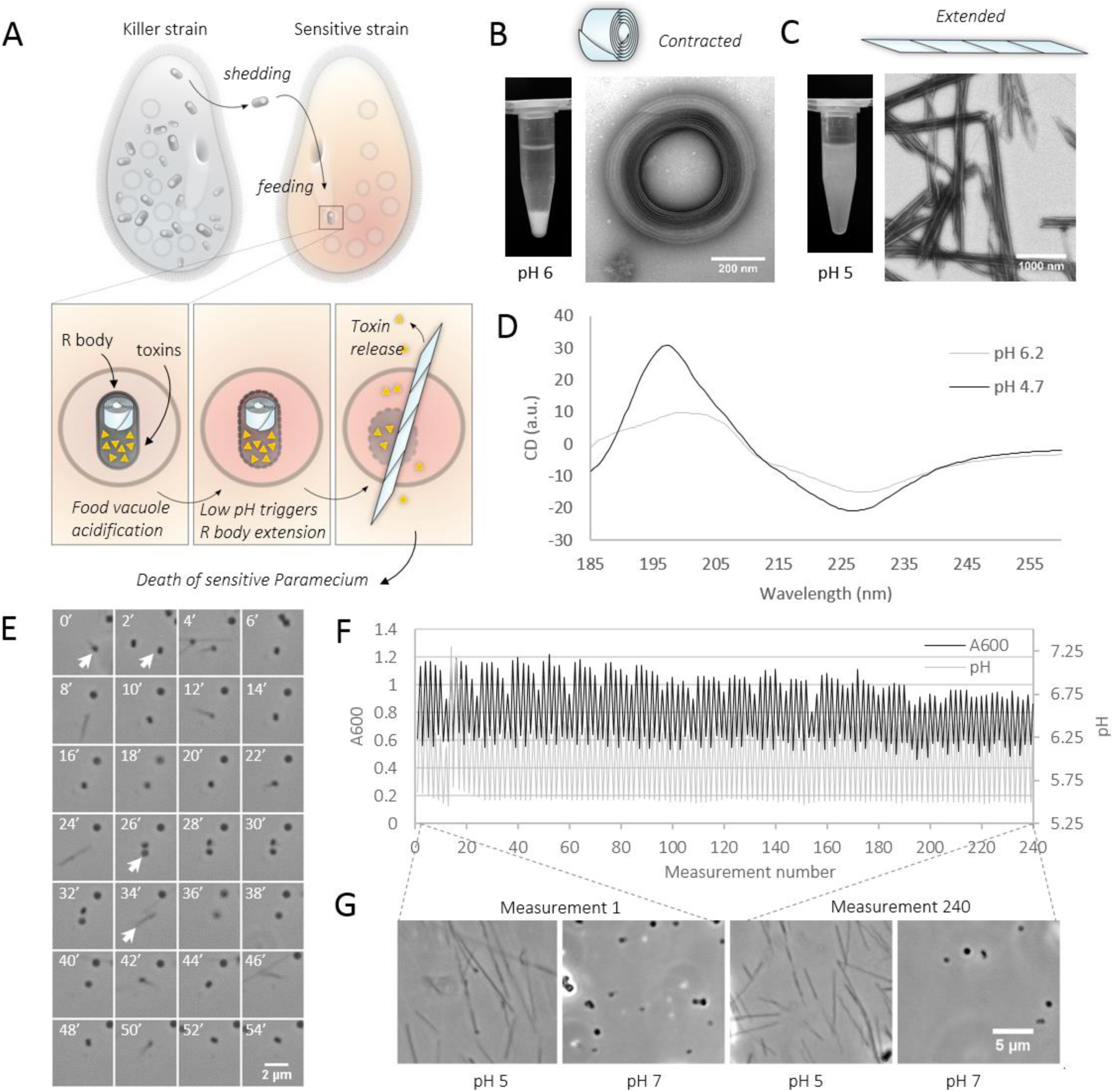
R bodies are reversible pH-driven pistons. A) R bodies are produced as cytoplasmic inclusion bodies by *C. taeniospiralis*, an endosymbiont of “killer strains” of *P. tetraurelia.* These bacteria are shed into the environment where they are consumed by “sensitive strains” of *Paramecium.* When the food vacuole acidifies, the R body extends to rupture the food vacuole, theoretically delivering toxins to the cytoplasm of the sensitive paramecium. B) At pH >5.7, R bodies are contracted, and they sediment. C) At pH <5.7, R bodies are extended, and remain in solution. D) These states produce different circular dichroism spectra. E) Montage (2 minute intervals) of purified R bodies imaged through multiple buffer changes. A single R body (white arrow) can be seen to undergo 9 cycles of extension and contraction. F) A bulk solution of R bodies was exposed to 240 pH changes (grey line), which produces absorbance changes (black line). G) Samples from the first and last solution changes are indistinguishable after each is resuspended in pH 5 and 7 solutions.

Type 51 R bodies can revert to their contracted form when the pH is raised (10), and they are resistant to harsh conditions including salt, detergents, and heat (11). These structures can be expressed in *E. coli* (7, 11, 8) from an operon of four open reading frames, *rebA-D*, two of which (RebA and RebB) are major structural proteins (12).

The process of R body extension is a simple, brute-force solution to the challenge of endosomal escape, and it therefore may hold promise as a research or clinical tool. However, it is not known whether R bodies can puncture membranes outside their natural setting, and this process has yet to be dynamically observed in any context. Furthermore, the mechanism by which R bodies convert chemical energy from protonation to protrusive forces is not understood, and it is unclear if they can extend identically multiple times.

Here we report that R bodies are able to undergo many cycles of extension and contraction *in vitro.* We also describe a simple assay that enables quantitation of the pH response of R bodies and its application in a screen for mutant R bodies that switch conformations at lower pH. Informed by this screen, we designed additional mutants that switch conformations at higher pH. Finally, we demonstrate that R bodies can release cytoplasmic contents from *E. coli* by rupturing the cell membrane.

## Results

### Production of functional R bodies in E. coli

We expressed the *reb* locus (13) in *E. coli* cells and purified R bodies based on their ability to sediment. R bodies from our *E. coli* expression system (Materials and Methods) behave as predicted (7, 10): at high pH, they resemble coils of ribbon (Figure 1B) by negative stain transmission electron microscopy. At low pH, they instead form extended, hollow tubes with pointed ends (Figure 1C). While R bodies were constantly 500nm in diameter at high pH and 165nm in diameter at low pH, we noted that the width of the ribbon appeared to vary between 100-600nm (Supplemental Figure 1). This distribution is broader than the 400nm ribbon width reported for natural R bodies (reviewed in (1)), but not unexpected given the presence of a variety of ribbon widths in previous reports of ectopically expressed R bodies (8).

### R body extension results in secondary structure changes and a macroscopic phenotype

R bodies display two distinct circular dichroism spectra at high and low pH (Figure 1D, Supplemental Figure 2). Analysis of this data suggests that R bodies are dominated by helical secondary structure, slightly more so at low pH than at high pH (Supplemental Figure 2).

We also found that R body solubility depends on their extension state. R bodies in the contracted, high pH state will sediment after several hours at room temperature, while those in the extended, low pH state remain in solution (Figure 1B and C). This difference can be rapidly appreciated with the naked eye in tubes or in 96-well plates.

### R bodies can undergo many cycles of extension and contraction

R bodies are capable of undergoing many dozens of cycles of extension and contraction without any apparent loss of function. The ability of R bodies to revert to their contracted state had been previously reported (10), but the limits of this reversibility remain an open question. Using phase contrast microscopy and flow cells that permitted on-stage buffer changes, we observed that single R bodies also undergo multiple cycles of extension and contraction in response to pH modulation (Figure 1E). A full-frame movie of this process is provided in Supplemental Movie 1.

To further probe the limits of this reversibility, we sequentially altered the pH of a solution of R bodies for 120 cycles (Figure 1F), reserving aliquots of R bodies at each step to confirm their extension state. Using phase contrast microscopy, we found that the first and last aliquots showed no detectable difference in their ability to respond to either high or low pH (pH 7.0 and 5.0, respectively) as measured by phase contrast microscopy (Figure 1G).

### Measuring R body extension in varied ionic strength solutions with a high-throughput assay

We used the differential solubility of extended and contracted R bodies described above to develop a rapid, quantitative, and high-throughput assay for R body extension state. In a 96 well plate reader, the pellets formed by contracted R bodies create a higher absorbance value than the same concentration of R bodies in solution. This difference can be further enhanced by programming the plate reader to agitate the plate before the reading; this reproducibly focuses the material toward the center of the well (Figure 2A). In this fashion, R body state can be measured across many conditions in a short period of time. We first used this method to confirm that the sequential pH changes described above indeed cause R bodies to change state (Figure 1F).

**Figure 2:**
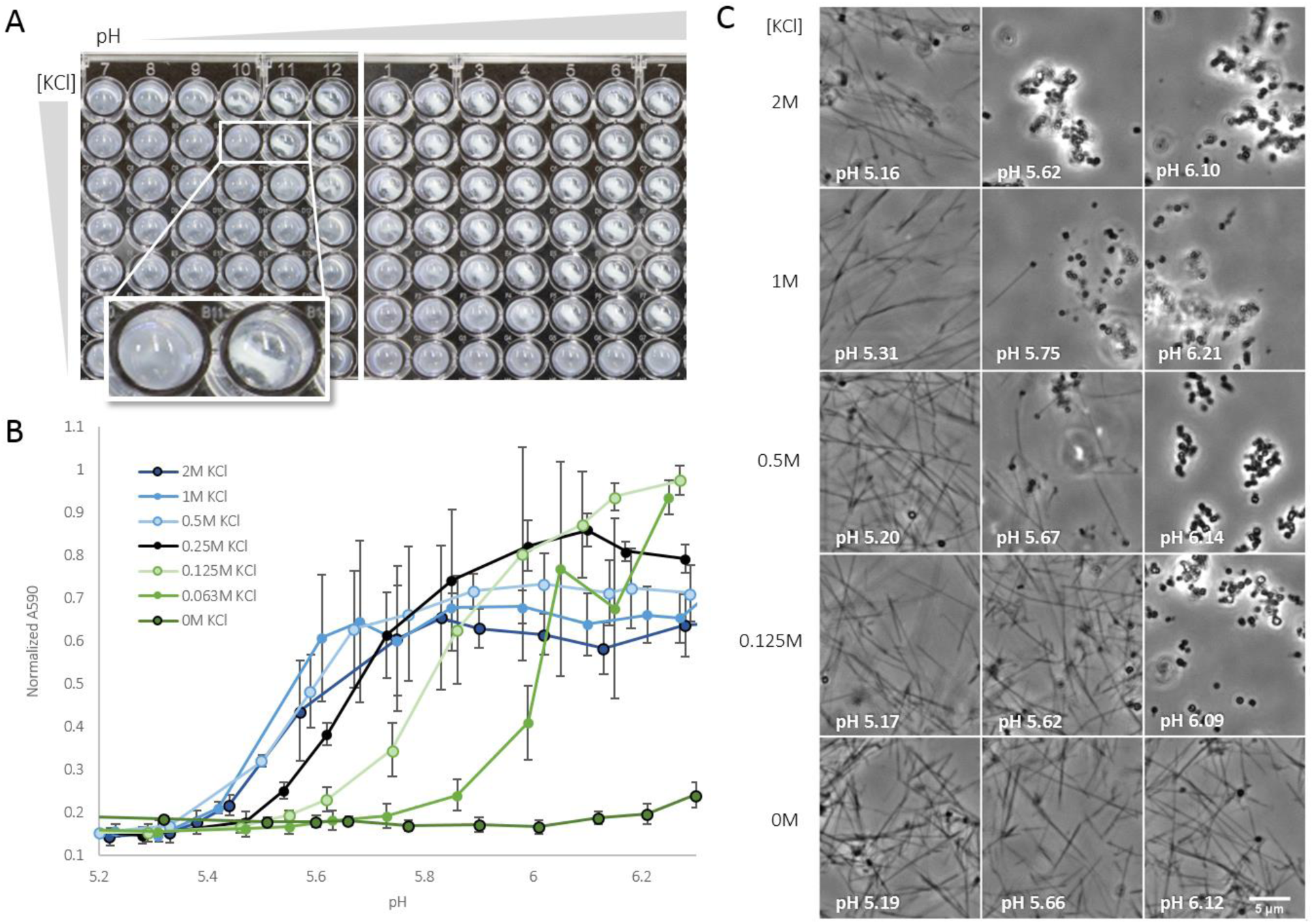
A high-throughput spectrophotometric assay for R body extension reveals dependence on ionic strength. A) 96-well plates with pH and KCl titration of R bodies. Lines in wells to the upper right are sedimented R bodies that have been “focused” by the shaking of the plate reader. B) Quantitation of (A) normalized to minimum A590 with standard deviation error bars; n=3. C) Phase contrast images of selected conditions in A and B.

Using this assay, we studied the impact of ionic strength on R body extension state. At high salt concentrations, the highest pH at which R bodies are completely extended is approximately 5.4. Meanwhile, at very low ionic strength, R bodies remain completely extended above pH 6.2. These data also reveal that within any given series, absorbance increases with pH, creating a sigmoidal curve with a median value that we refer to as the conversion pH (Figure 2B). We verified that the observed transfer curve reflects actual morphological changes in R bodies by visualizing their state with phase contrast microscopy (Figure 2C).

### A screen for R bodies that extend at variable pHs identifies changes in a region of RebA

We adapted our plate-based assay as a screening tool for R bodies defective in their pH response. We amplified a region spanning the *rebA* and *rebB* open reading frames with error-prone PCR and cloned this region into a plasmid backbone containing an unmodified copy of the remainder of the *reb* operon, *rebC* and *rebD* (Figure 3A). Single colonies resulting from a transformation of this library into C43 *E. coli* cells were then selected for growth in a 96-well plate format. R bodies were expressed and purified in transferred to an optically-clear plate at pH 5.5. Under these conditions, wild type R bodies will remain soluble, but mutants that require a lower pH to extend will sediment (Figure 3A, right panel). We therefore visually selected wells with a dense, visible pellet (like those highlighted in Figure 3B) as putative hits. These were subsequently confirmed by pH titration (Figure 3C) and classified according to their approximate conversion pH.

**Figure 3:**
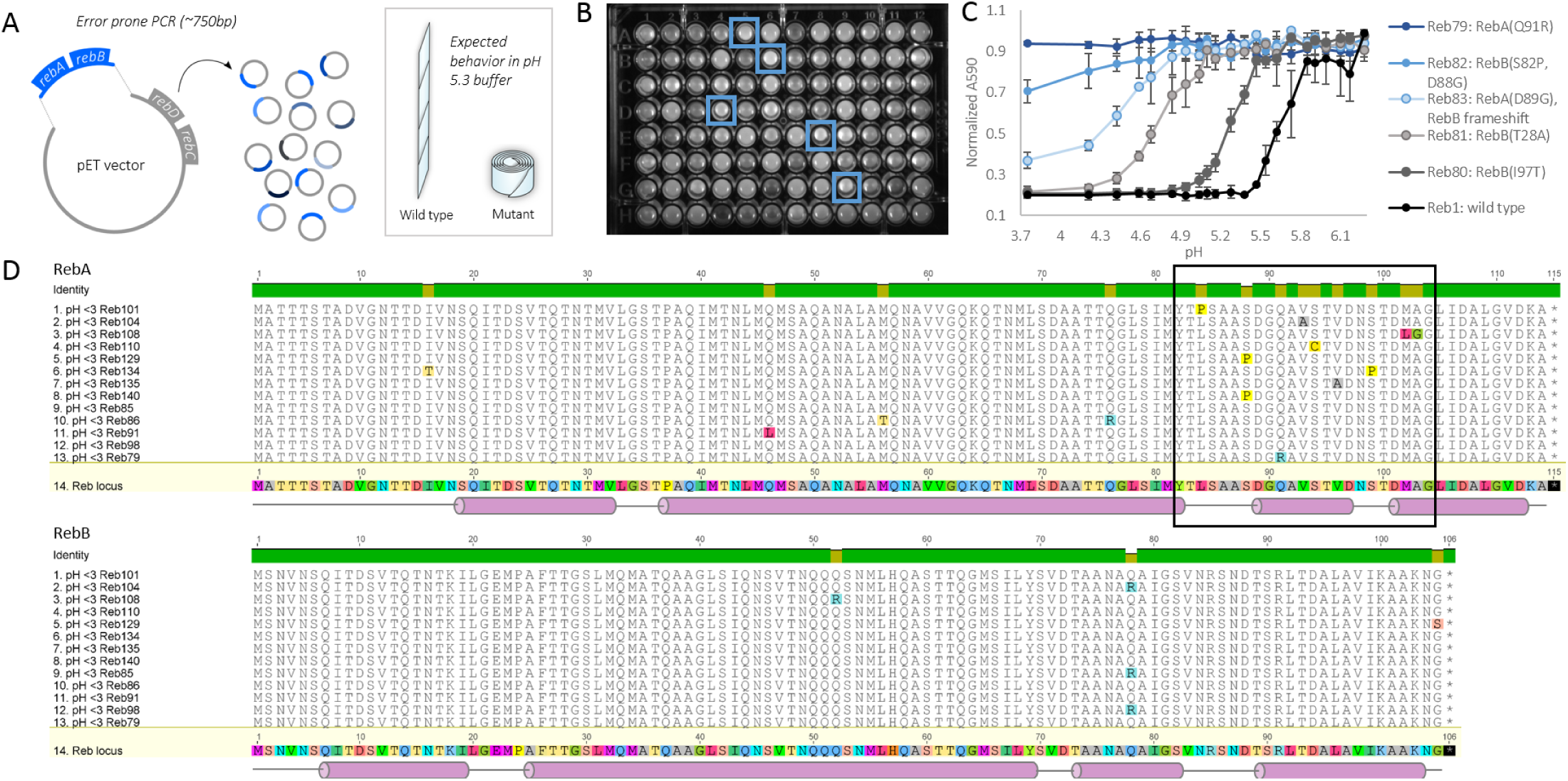
A visual screen for mutants defective in pH response identifies a region in RebA that mediates extension. A) Design of the screen. RebA and B were amplified with error-prone PCR and ligated into a backbone containing the remainder of the *reb* operon to create a library of mutants that were expressed and purified in a 96-well format. The R bodies were resuspended in pH 5.5, at which wild type R bodies would be extended, but mutants would be contracted. B) Mutants that were sedimented were visually selected. C) Absorbance series of selected mutants showing the range of phenotypes isolated, normalized to maximum A590 with standard deviation error bars; n=3. D) Amino acid alignment of the mutants unable to extend at pH 3.0. Secondary structure predictions of the wild-type sequence by PSI-PRED (cylinders = helix, line = coil). Highlighted region shows clustering in the C-terminal region of RebA.

Out of a library of 1728 clones, we identified 60 isolates defective in pH response (representative isolates shown in figure 3C, sequences shown in Supplemental figures 3 and 4). Of these, 13 did not extend below pH 3.0 (Figure 3D), though mutants from this class resemble normally assembled R bodies by negative stain electron microscopy (Supplemental figure 5). These clones possessed a total of 16 unique mutations, 9 of which fall into a 20 base pair region at the C-terminus of RebA. This region accounts for under 10% of the amino acids covered in the error-prone PCR reaction, but contains over 50% of the mutations that result in severely defective R bodies, leading us to hypothesize that it plays a role in the extension process. Three of the unique mutations in this region replace residues with proline.

### Rational design produces mutants with increased pH sensitivity

Introducing alanine residues in the region identified by the screen generated R bodies with higher conversion pH. To probe the effect of residues classically thought to stabilize or destabilize helices (14), we constructed a series of mutants with either individual residues or tracts of residues replaced with alanines or prolines (Figure 4A). We identified three mutants that enable R bodies to extend at higher pH than wild type, with the most dramatic being RebA S88A (Figure 4B). This phenotype is also evident by phase contrast microscopy (Figure 4C).

**Figure 4:**
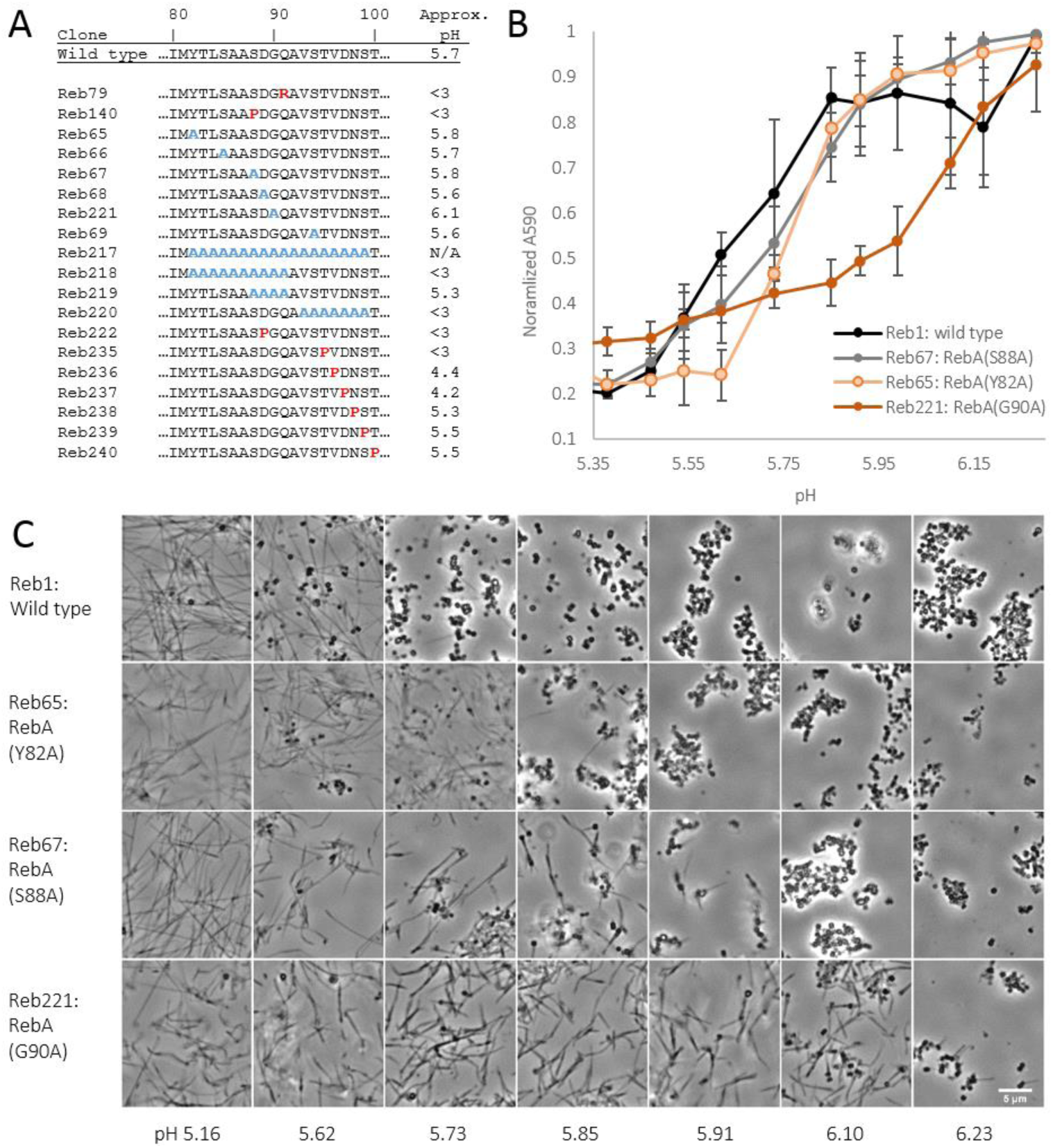
Rational design of gain of function mutants. A) List of mutants affecting the C-terminal region of RebA. Reb79 and 140 were identified in the previously-described screen (Figure 3) while the others were made deliberately. B) Quantitation of the behavior of mutants isolated in the screen with a higher conversion pH than wild type, normalized to maximum A590 with standard deviation error bars; n=3. C) Phase contrast imaging of the mutants in (B).

### R bodies are capable of rupturing *E. coli* spheroplasts

R bodies can act as membrane-breaking devices outside of their natural context. We constructed a plasmid encoding functional fluorescent R bodies as well as a soluble fluorescent protein, mCherry (Materials and Methods). After expressing this construct in *E. coli*, we treated cells with lysozyme and imaged them in flow chambers through to which we added low pH buffer that contained salts (methylamine hydrochloride and potassium benzoate) shown to disrupt *E. coli’s* otherwise robust pH homeostasis (15) (Figure 5A). When spheroplasts containing functional (wild type) R bodies were exposed to low pH buffer, only a minority remained intact (36%, Figure 5B). The loss of fluorescence was often accompanied by dramatic protrusions of R bodies that distend cells (Figure 5E and G, right side of Supplemental Movie 2). R bodies sometimes extended inside cells before the membrane ruptured (Figure 5G, leftmost 4 cells), suggesting that extension causes lysis. By contrast, when cells from the same culture were not treated with lysozyme, most (95%) remained intact, as did spheroplasts not treated with salts to disrupt pH homeostasis (89%) and spheroplasts cells containing R bodies with the RebA S88P mutation, which are incapable of extension (90%) (Figure 5B-F, left side of Supplemental Movie 2). Thus, R body extension, rather than the presence of R bodies or the stress of their assembly, lyses cells at low pH.

**Figure 5:**
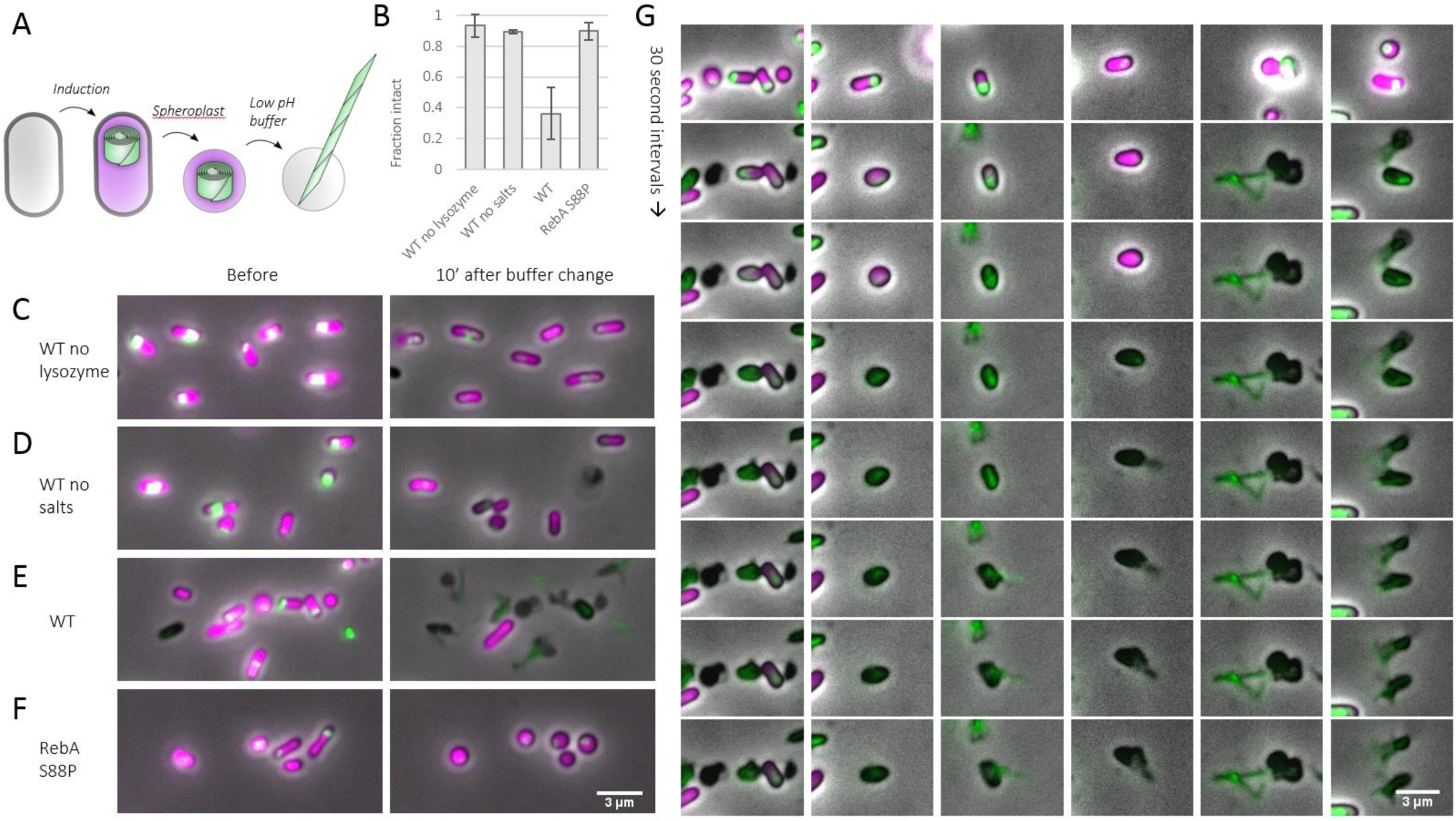
R body extension is capable of breaking membranes *in vitro.* A) *E. coli* cells were induced to produce mNeon-labeled R bodies and soluble mCherry. These cells were then spheroplasted to remove the cell wall and treated with salts (methylamine hydrochloride and potassium benzoate) to destroy proton homeostasis in conjunction with a buffer change. Cells were imaged in flow cells by fluorescence microscopy to record their behavior. B) Fraction of cells still mCherry-positive 10 minutes after buffer change. Measured from three independent experiments with >50 cells per experimental condition. Error bars: standard deviation of each experiment’s average. C-F) Representative images before (left) and 10 minutes after (right) replacement of buffer with 100mM MES pH 4.9, 40mM potassium benzoate, and 40mM methylamine hydrochloride. C) Functional R bodies in cells not spheroplasted. D) Functional R bodies in 100mM MES pH 4.9 without salts (no potassium benzoate or methylamine hydrochloride). E) Functional R bodies with salts. F) Non-functional R bodies (RebA S88A) with salts. G) Montages of individual wild type R body lysis events. Time interval is 30 seconds. Left-most image shows same cells as in panel E.

## Discussion

### R bodies as reversible protein machines

Our data show that type 51 R bodies are capable of multiple rounds of extension and contraction, suggesting that all of the energy for their transformations comes from chemical changes in the buffer instead of from other sources such as protein folding. This distinguishes type 51 R bodies from many other protrusive apparatuses in biology such as acrosomes, trichocysts, and nematocysts. Instead, they are behaviorally similar to other polymeric structures like forisomes (16) and spasmonemes (17), which are driven by changes in Ca^2+^ concentration to switch between extended and contracted states.

### Mechanism of action

We speculate that R body extension requires the formation or extension of a helix in RebA. Such rearrangements would have precedent in the loop-to-helix transition that drives the pH-dependent rearrangement of viral hemagglutinin (18). Analysis of our R body circular dichroism data (Figure 1D) suggests a slightly greater contribution of helices to the low pH spectrum than to the high pH spectrum (Supplemental Figure 2). Though this difference accounts for only 2% of the residues, these may bridge the small, unstructured regions predicted in the C-terminal region of RebA (PSIPRED predictions shown in Figure 3D). Many of the mutations identified in the screen introduce proline residues, which can disrupt helices (14). Therefore, these mutations may prevent the pH-dependent formation of a helix in this C-terminal region. Conversely, alanine residues can stabilize helices, so it is unsurprising that some of the rationally designed alanine mutants we produced bias R bodies toward an extended conformation.

We have shown that when R bodies are in buffers of high ionic strength, a lower pH is required to contract them than is needed at low ionic strength (Figure 2B and C). As K^+^ and Cl^−^ fall relatively early in the Hofmeister series, high concentrations of salt may function to increase surface tension and therefore strengthen interactions between hydrophobic residues. These hydrophobic interactions may play a role in R body contraction.

### Implications for bioengineering

Because R bodies are robust to buffer changes and capable of functioning in a cell-independent fashion, they should be functional in a wide variety of biotechnology applications. The ability to engineer R body pH sensitivity expands their potential utility in a range of diverse contexts, especially in delivering molecules across biological barriers. For example, in an application that parallels their proposed natural function, R bodies could be used to enhance endosomal delivery of DNA, RNA, or other bioactive molecules. While the pH of most late endosomes is below 5.7, some cell types do not severely acidify the contents of their phagosomes (19), and in this case, R bodies engineered with increased pH sensitivity (such as the RebA S88A mutant) would be advantageous. These engineered R bodies could also be used in other scenarios with mild pH conditions. For example, they may be able to specifically occlude or puncture tumor microvasculature, where hypoxic conditions cause a drop in blood pH.

Type 51 R bodies are just one among several described types of R bodies, all of which have been reported to have varied properties (reviewed in (1)). Given that genes encoding R body homologs have recently been identified in a wide variety of bacteria (20), we may be able to produce actuators of different sensitivities and strengths by harnessing this natural diversity.

## Materials and Methods

### Cloning

The Reb locus (13) was Gibson assembled into pETM11 using gBlocks from Integrated DNA Technology under the control of the vector’s T7 promotor. Site directed mutagenesis was performed with New England Biolab’s Q5 kit. To screen for pH variant R bodies, the RebA and RebB coding sequences were amplified with error-prone PCR (EP-PCR) using Taq polymerase in the presence of 62.5 and 125uM MnCl2 to introduce errors. This product was then cloned into an unmutated backbone containing the remainder of the *reb* operon. To make fluorescent R bodies, we fused mNeonGreen (21) to the N-terminus of a second copy of RebB under the control of a weak RBS, BBa_B0033 from the Registry of Standard Biological Parts. This was cloned downstream of an mCherry ORF with a strong RBS, which itself was cloned downstream of the other ORFs in the operon. A list of constructs used in this study is available in Supplementary Table 1.

### R body expression and purification

Plasmids containing R bodies were transformed into C43 cells (22), a derivative of BL21 with attenuated T7 expression (23). Cells were grown in TPM to OD 600 0.2-0.6 and induced with 1mM IPTG. Expression proceeded for 18 hours at 37° C.

To purify R bodies, cell pellets were flash-frozen in liquid nitrogen, then thawed. Cells were resuspended in 25mM Tris pH 7.5, 100mM NaCl, and 2mM EDTA. Egg white lysozyme was added to a concentration of approximately 17 µg/ml, and cells were incubated at 37° C for 1 hour. After this time, the buffer was adjusted to contain 10mM MgCl2, 10mM CaCl2, and approximately 15 µ/ml DNAse from bovine pancreas. Cells were once again incubated at 37° C for 20 minutes. Next, the buffer was adjusted to contain 1% SDS. After manual mixing, cells were spun at 4,000 RPM in a tabletop centrifuge for 20 minutes to pellet the R bodies. The R body pellet was then washed three times by resuspension in water, followed by spins, as above.

R bodies were stored at 4° C for short term use (<1 week), or at −80° C after flash-freezing in liquid nitrogen in the presence of 25mM Tris pH 7.5, 100mM KCl, and 15% glycerol.

### Electron microscopy

200 mesh formvar and carbon-coated copper grids (Electron Microscopy Sciences) were glow discharged for 30 seconds before the application of R bodies. These were washed by applying the grids sequentially to three drops of buffer and three drops of either 0.75% uranyl formate or 1% uranyl acetate, wicking with filter paper between each wash. Grids were visualized on either a JEOL 1200EX or a Tecnai G2 Spirit BioTWIN.

### Circular dichroism

R bodies at a concentration of 0.2mg/ml (calculated by Bradford) were washed into 0.1M sodium phosphate buffer at pH 6.2 and 4.7. Data was collected on a JASCO J-815 Circular Dichroism Spectropolarimeter. Data was analyzed with DichroWeb (24, 25) predictions based on the CDSSTR method (26, 27) and reference set 3 (28).

### Light microscopy

Flow cells used in spheroplast and kinetics experiments were constructed from a 22x22mm coverslip adhered to a 22×60mm coverslip with Scotch double-sided tape. For kinetics experiments, coverslips were heated at 50° for 4 hours in 1M HCl, then washed and sonicated in water, then ethanol for 30 minutes each. Completed flow cells were then incubated for 5 minutes in 10μl of PBS containing 1mg/ml BSA (1/20^th^ of which was labeled with biotin). After washing with PBS, cells were incubated with 25μg/ml streptavidin and washed again. At this point, wild type R bodies that had been nonspecifically labeled with maleimide-biotin (15 minutes at room temperature with 1mM EZ-link maleimide-PEG2-biotin, Thermo Fisher Scientific) were added and left to incubate for 5 minutes. The chambers were washed with more PBS followed by 1mg/ml BSA/BSA-biotin solution. Finally, during imaging, pH was changed by flowing 30-50μl of citric acid-Na_2_HPO_4_ buffer at either pH 5.2 or pH 6.2 through the flow cell.

Static images of purified R bodies were obtained from simple wet mounts on untreated coverslips and slides.

All images were acquired on a Nikon TE2000 microscope equipped with a 100x phase objective, Perfect Focus, and an Orca ER camera.

### Spectrophotometric assay

The absorbance of R bodies (100μl in each well of a 96-well plate) at 600nm or 590nm was read on a Perkin Elmer Victor^3^V plate reader following 15 seconds of agitation by the plate reader.

### Screen for mutants defective in pH response

The error-prone PCR-generated library described above (Cloning) was transformed into *E. coli* C43 cells, and single colonies were picked to 1ml of TPM in 96-well assay blocks. R bodies were expressed and purified as described above. After washing R bodies in water sequentially, they were resuspended in 250mM MES pH 5.5 and 250mM KCl. Hits were visually identified by the sedimentation of R bodies in the respective well and confirmed by measuring the behavior against a pH series as in Figure 2C.

### Sequence analysis

Sequences were aligned with Geneious version 8.1 (Biomatters Ltd). Secondary structure prediction was done with PSIPRED (29, 30).

### Spheroplasting

Because R body expression takes several hours, we employed a spheroplasting method described previously that works even in stationary phase (31). Briefly, cells expressing R bodies that had been growing at 37° for 4-12 hours after induction were harvested and washed in 200mM Tris pH 8.0, then resuspended in the same buffer. This was diluted 1:1 with 200mM Tris pH 8.0 containing 1M sucrose and 1mM EDTA. 10ul of a solution containing 7 mg/ml lysozyme was added, and the mixture was incubated at room temperature for 20 minutes. The buffer was adjusted to 20mM MgCl_2_ and cells were loaded into flow cells (see Microscopy). The flow cell was washed with buffer containing 200mM Tris pH 8.0, 0.5M sucrose, and 0.5mM EDTA prior to the start of imaging. The buffer was then swapped to contain 100mM MES pH 4.9 with or without 40mM methylamine hydrochloride and 40mM potassium benzoate, a combination that has been previously used to destroy *E. coli’s* pH homeostasis (15).

## Acknowledgements

We are grateful to Timothy Mitchison (HMS), Michael Baym (HMS), Ethan Garner (Harvard), Justin Kollman (UW), Rob Phillips (Caltech), Dan Fletcher (UC Berkeley) and Michael Vahey (UC Berkeley) for helpful discussions. We are grateful to Genevieve Dobihal (Harvard), Nathan Rollins (Harvard), and Brendan Cruz (Harvard) for constructive feedback on the manuscript.This work was supported by a Jane Coffin Childs Postdoctoral Fellowship to JKP and an Office of Naval Research MURI Grant N00014-11-1-0725.

## References

1. Pond FR, Gibson I, Lalucat J, Quackenbush RL. 1989. R-body-producing bacteria. Microbiol Rev 53:25–67.

2. Mueller JA. 1965. Vitally stained kappa in Paramecium aurelia. J Exp Zool 160:369–372.

3. Sonneborn TM, Jacobson W, Dippell RV. 1946. Paramecin 51, an antibiotic produced by Paramecium aurelia; amounts released from killers and taken up by sensitives; conditions protecting sensitives. Anat Rec 96:514.

4. Austin ML. 1946. Contributions towards an analysis of the killing action of variety 4 killers in Paramecium aurelia. Anat Rec 96:514.

5. Jurand A, Rudman BM, Preer JR. 1971. Prelethal effects of killing action by stock 7 of Paramecium aurelia. J Exp Zool 177:365–387.

6. Preer LB, Jurand A, Preer JR, Rudman BM. 1972. The Classes of Kappa in Paramecium Aurelia. J Cell Sci 11:581–600.

7. Quackenbush RL, Burbach JA. 1983. Cloning and expression of DNA sequences associated with the killer trait of Paramecium tetraurelia stock 47. Proc Natl Acad Sci U S A 80:250–254.

8. Schrallhammer M, Galati S, Altenbuchner J, Schweikert M, Görtz H-D, Petroni G. 2012. Tracing the role of R-bodies in the killer trait: Absence of toxicity of R-body producing recombinant E. coli on paramecia. Eur J Protistol 48:290–296.

9. Kusch J, Czubatinski L, Wegmann S, Hubner M, Alter M, Albrecht P. 2002. Competitive advantages of Caedibacter-infected Paramecia. Protist 153:47–58.

10. Preer Jr. JR, Hufnagel LA, Preer LB. 1966. Structure and behavior of R bodies from killer paramecia. J Ultrastruct Res 15:131–143.

11. Kanabrocki JA, Quackenbush RL, Pond FR. 1986. Organization and expression of genetic determinants for synthesis and assembly of type 51 R bodies. J Bacteriol 168:40–48.

12. Heruth DP, Pond FR, Dilts JA, Quackenbush RL. 1994. Characterization of genetic determinants for R body synthesis and assembly in Caedibacter taeniospiralis 47 and 116. J Bacteriol 176:3559–3567.

13. Jeblick J, Kusch J. 2005. Sequence, Transcription Activity, and Evolutionary Origin of the R-BodyCoding Plasmid pKAP298 from the Intracellular Parasitic BacteriumCaedibacter taeniospiralis. J Mol Evol 60:164–173.

14. Pace CN, Scholtz JM. 1998. A helix propensity scale based on experimental studies of peptides and proteins. Biophys J 75:422–427.

15. Martinez KA, Kitko RD, Mershon JP, Adcox HE, Malek KA, Berkmen MB, Slonczewski JL. 2012. Cytoplasmic pH response to acid stress in individual cells of Escherichia coli and Bacillus subtilis observed by fluorescence ratio imaging microscopy. Appl Environ Microbiol 78:3706–3714.

16. Knoblauch M, Noll GA, Müller T, Prüfer D, Schneider-Hüther I, Scharner D, van Bel AJE, Peters WS. 2003. ATP-independent contractile proteins from plants. Nat Mater 2:600–603.

17. Upadhyaya A, Baraban M, Wong J, Matsudaira P, van Oudenaarden A, Mahadevan L. 2008. Power-Limited Contraction Dynamics of Vorticella convallaria: An Ultrafast Biological Spring. Biophys J 94:265–272.

18. Harrison SC. 2008. Viral membrane fusion. Nat Struct Mol Biol 15:690–698.

19. Canton J, Khezri R, Glogauer M, Grinstein S. 2014. Contrasting phagosome pH regulation and maturation in human M1 and M2 macrophages. Mol Biol Cell 25:3330–3341.

20. Raymann K, Bobay L-M, Doak TG, Lynch M, Gribaldo S. 2013. A Genomic Survey of Reb Homologs Suggests Widespread Occurrence of R-Bodies in Proteobacteria. G3 GenesGenomesGenetics 3:505–516.

21. Shaner NC, Lambert GG, Chammas A, Ni Y, Cranfill PJ, Baird MA, Sell BR, Allen JR, Day RN, Israelsson M, Davidson MW, Wang J. 2013. A bright monomeric green fluorescent protein derived from Branchiostoma lanceolatum. Nat Methods 10:407–409.

22. Miroux B, Walker JE. 1996. Over-production of proteins in Escherichia coli: mutant hosts that allow synthesis of some membrane proteins and globular proteins at high levels. J Mol Biol 260:289–298.

23. Wagner S, Klepsch MM, Schlegel S, Appel A, Draheim R, Tarry M, Högbom M, van Wijk KJ, Slotboom DJ, Persson JO, de Gier J-W. 2008. Tuning Escherichia coli for membrane protein overexpression. Proc Natl Acad Sci U S A 105:14371–14376.

24. Whitmore L, Wallace BA. 2004. DICHROWEB, an online server for protein secondary structure analyses from circular dichroism spectroscopic data. Nucleic Acids Res 32:W668–W673.

25. Whitmore L, Wallace BA. 2008. Protein secondary structure analyses from circular dichroism spectroscopy: Methods and reference databases. Biopolymers 89:392–400.

26. Compton LA, Johnson WC. 1986. Analysis of protein circular dichroism spectra for secondary structure using a simple matrix multiplication. Anal Biochem 155:155–167.

27. Manavalan P, Johnson Jr. WC. 1987. Variable selection method improves the prediction of protein secondary structure from circular dichroism spectra. Anal Biochem 167:76–85.

28. Sreerama N, Woody RW. 2000. Estimation of Protein Secondary Structure from Circular Dichroism Spectra: Comparison of CONTIN, SELCON, and CDSSTR Methods with an Expanded Reference Set. Anal Biochem 287:252–260.

29. Jones DT. 1999. Protein secondary structure prediction based on position-specific scoring matrices. J Mol Biol 292:195–202.

30. Buchan DWA, Minneci F, Nugent TCO, Bryson K, Jones DT. 2013. Scalable web services for the PSIPRED Protein Analysis Workbench. Nucleic Acids Res 41:W349–357.

31. Witholt B, Boekhout M, Brock M, Kingma J, Heerikhuizen HV, Leij LD. 1976. An efficient and reproducible procedure for the formation of spheroplasts from variously grown Escherichia coli. Anal Biochem 74:160–170.

